# An in vivo biomarker to characterize ototoxic compounds and novel protective therapeutics

**DOI:** 10.1101/2022.05.31.493913

**Authors:** Joseph A. Bellairs, Van A. Redila, Patricia Wu, Ling Tong, Alyssa Webster, Julian A. Simon, Edwin W. Rubel, David W. Raible

## Abstract

There are no approved therapeutics for the prevention of hearing loss and vestibular dysfunction from drugs like aminoglycoside antibiotics. While the mechanisms underlying aminoglycoside ototoxicity remain unresolved, there is considerable evidence that aminoglycosides enter inner ear mechanosensory hair cells through the mechanoelectrical transduction (MET) channel. Inhibition of MET-dependent uptake with small molecules or modified aminoglycosides is a promising otoprotective strategy. To better characterize mammalian ototoxicity and aid in the translation of emerging therapeutics, a biomarker is needed. In the present study we propose that neonatal mice systemically injected with the aminoglycosides G418 conjugated to Texas Red (G418-TR) can be used as a histologic biomarker to characterize in vivo aminoglycoside toxicity. We demonstrate that postnatal day 5 mice, like older mice with functional hearing, show uptake and retention of G418-TR in cochlear hair cells following systemic injection. When we compare G418-TR uptake in other tissues, we find that kidney proximal tubule cells show similar retention. Using ORC-13661, an investigational hearing protection drug, we demonstrate *in vivo* inhibition of aminoglycoside uptake in mammalian hair cells. This work establishes how systemically administered fluorescently labeled ototoxins in the neonatal mouse can reveal important details about ototoxic drugs and protective therapeutics.

## 1 Introduction

Aminoglycosides are potent, broad-spectrum antibiotics that are clinically employed to treat mycobacterial infections (*mycobacterium tuberculosis)*, Gram-negative urinary and respiratory infections, and severe neonatal infections (Durante-Mangoni et al., 2009). Despite their wide antibacterial coverage, low cost, and stability at ambient temperatures, use of aminoglycosides in developed countries is generally limited by dose-dependent damage to the kidney (nephrotoxicity) and inner ear (ototoxicity) (Forge and Schacht, 2000; Rizzi and Hirose, 2007; Van Boeckel et al., 2014). Aminoglycoside nephrotoxicity can be monitored for with regular kidney function screening and injury can be minimized or even reversed with strategies like adjusting dose for kidney function, hydration, limiting therapy duration, and once-daily dosing (Munckhof et al., 1996; Hatala et al., 1996; Bailey et al., 1997; Guo and Nzerue, 2002). To date there are no reliable clinical strategies or therapeutics that are approved or used to prevent or minimize aminoglycoside ototoxicity.

Aminoglycoside ototoxicity targets the mechanosensory hair cells within the inner ear, which convert mechanical energy from sound waves into electrical impulses. These cells are also exceptionally vulnerable to a wide variety of genetic and environmental challenges, including excessive noise exposure and off-target effects from therapeutics such antineoplastic agents in addition to aminoglycoside antibiotics (Forge and Schacht, 2000; Rizzi and Hirose, 2007). While aminoglycoside antibiotics have been known to cause disabling hearing loss and vestibular disfunction for decades (Rybak et al., 2008), the cellular mechanisms underlying this pathology remain under intensive study. Previous work has implicated a diverse array of cellular insults caused by aminoglycosides, including excessive free radical accumulation (Priuska and Schacht, 1995; Hirose et al., 1997), mitochondrial damage (Qian and Guan, 2009), plasma-membrane interactions (Marche et al., 1983), calcium dysregulation (Esterberg et al., 2014, 2016), and both caspase-dependent (Cheng et al., 2003; Coffin et al., 2013) and caspase-independent (Jiang et al., 2006) cell death pathways. The diverse range of cytotoxic effects attributed to aminoglycosides makes target selection and rational drug design for protective therapeutics challenging. As an alternative approach, previous studies employed drug and compound screens for phenotypic drug discovery using the zebrafish model. Zebrafish lateral line hair cells closely resemble mammalian hair cells, and importantly they demonstrate susceptibility to known ototoxins such as aminoglycosides and the anticancer therapeutic cisplatin (Pickett and Raible, 2019). Utilizing high-throughput chemical screens in zebrafish to assess for hair cell protection from aminoglycosides, several promising protective compounds have been discovered (Owens et al., 2008a; Kenyon et al., 2021), optimized (Chowdhury et al., 2018), and advanced to clinical trials (Kitcher et al., 2019).

Despite recent successes, translation of preclinical otoprotective leads to mammals and ultimately human clinical trials remains an immense challenge. Indeed, across numerous biomedical fields a majority of drug discovery pipelines fail to traverse the preclinical and clinical divide (Seyhan, 2019). Many preclinical ototoxicity studies occur in zebrafish and/or mammalian neonatal explant models, and while these models faithfully recapitulate parts of the auditory system (e.g. hair cells), they lack the complex pharmacokinetics and pharmacodynamics of an in vivo mammalian model. In vivo mammalian studies assessing clinical outcomes such as hearing loss with pre- and post-treatment auditory brainstem response (ABR) thresholds are the gold standard, but these studies in mice and rats are resource-intensive and take weeks to months to complete. Translating in vitro exposure conditions to in vivo dosing regimens is also a complex endeavor when considering pharmacokinetics of multiple compounds and the assessment of efficacy is limited to the clinical endpoint of ABR threshold shifts. As such, in vivo studies are often reserved for only the most promising lead compounds, and even then, finding a safe and effective drug candidate can be likened to the Herculean task akin to finding a needle in a haystack.

A biomarker is a measured characteristic that is an indicator of a normal biologic process, a pathogenic process, or a response to an exposure or intervention (FDA-NIH Biomarker Working Group, 2016). For studies of ototoxicity, there is a clear need for rapid, low-cost predictive biomarkers that can determine the ability of interventions and therapeutics to inhibit the accumulation of ototoxins in mammalian hair cells in vivo. Previous studies have used a variety of preparations such as cochlear cultures from neonatal mice (Richardson and Russell, 1991) and derived cell lines such as HEI-OC1 cells (Kalinec et al., 2003), and more recent reports proposed circulating microRNAs (Lee et al., 2018) and proteins such as prestin (Naples et al., 2018) as specific markers of hair cell injury. However, the translational utility of these markers remain to be validated. Here, we propose utilizing in vivo hair cell uptake of fluorescently labeled aminoglycosides as a histologic biomarker for ototoxicity. Despite the variety of mechanisms described for aminoglycoside-induced ototoxicity, there is consensus that a fundamental property of susceptible cells is their uptake and retention of aminoglycosides. Previous studies demonstrated that proximal convoluted tubule cells within the kidney and inner ear mechanosensory hair cells show aminoglycoside uptake and retention following in vivo systemic aminoglycoside administration, and there appears to be good correlation between aminoglycoside uptake and cell death (Imamura and Adams, 2003; Dai et al., 2006; Kim and Ricci, 2022).

We examine intracellular accumulation of the aminoglycoside geneticin (G418) conjugated to the fluorophore Texas red (TR) in neonatal (P5) mouse hair cells following systemic injection. Building off recent studies (Makabe et al., 2020; Qian et al., 2021), we performed experiments in neonatal mice because (I) neonatal mice can be readily obtained in large number and at low costs from a small number of breeding adults and (II) neonatal otic capsules are incompletely ossified, allowing for rapid tissue fixation and easy dissection of sensory epithelium. We demonstrate that neonates, like juvenile mice (P25-30) with mature hearing sensitivity, exhibit dose- and time-dependent uptake and accumulation of G418-TR in hair cells. Using ORC-13661, an orally available small molecule that acts as a high-affinity permeant blocker of the mechanoelectrical transduction (MET) channel (Kitcher et al., 2019) and preserves hearing in rats treated with the aminoglycoside amikacin (Chowdhury et al., 2018), we demonstrate that hair cell uptake of G418-TR is inhibited in vivo in mammals. Our results show that neonatal mice injected with G418-TR can be used as a translational tool to better understand in vivo mammalian ototoxicity and characterize the efficacy of otoprotective therapeutics that block aminoglycoside uptake. These results also confirm a mechanism of action for ORC-13661 predicted from zebrafish and mammalian in vitro studies (Kitcher et al., 2019).

## 2 Materials and Methods

### 2.1 Animals

C57BL/6J wild-type mice were used in all experiments. Both male and female mice were used in all experiments. All procedures involving animals were carried out in accordance with the National Institutes of Health guide for care and use of laboratory animals and approved by the Institutional Animal Care and Use Committee of the University of Washington.

### 2.2 Synthesis of G418-TR

A modified synthesis of geneticin (G418) conjugated Texas Red (TR) was performed based on previously published protocols(Liu et al., 2015). TR-NHS ester was dissolved in anhydrous N,N-dimethylformamide and added to a solution of 1.377 mmol G418 disulfate in a 100 mM potassium carbonate solution. The resulting solution was mixed in darkness at 4°C for 48 h. G418-TR was purified by Reverse-phase chromatography connected to a UV-Vis detector monitoring a wavelength at 214nm. The analyte was gradient eluted from a Waters delta-pak C18 (300 angstrom) 19mmx 30cm column using buffers of 0.1% trifluoroacetic acid (by volume in water) for “A” and acetonitrile/0.08% trifluoroacetic acid (by volume) for “B”. Fractions were collected every thirty seconds between 15 and 38 minutes using a gradient as follows: 0 minutes at 5% “B”, followed by a 5-minute hold at 5% “B”, then ramping up to 20% “B” in one minute, followed by a linear gradient to 50% “B” in 25 minutes, followed by another linear ramp to 95% “B” in 2 minutes and holding at 95% “B” for 5 minutes, and finally ramping down to 5% “B” in 2 minutes and column equilibration at 5% “B” over another 5 to 10 minutes. Up to 1/3 of the sample volume was loaded onto the column and purified in three successive HPLC runs. Those fractions in which the m/z value corresponding to the analyte of interest were pooled, lyophilized, and stored in the dark at −20°C with a final yield of 68%.

### 2.3 G418-TR uptake

G418, an aminoglycoside structurally related to gentamicin, was used for all experiments because unlike gentamicin, which is supplied as a variable mixture of aminoglycosides, G418 is a purified single aminoglycoside without batch-to-batch variation (O’Sullivan et al., 2020). G418-TR has previously been used to study aminoglycoside trafficking in zebrafish hair cells and it has been shown to be similar to gentamicin, both in toxicity and intracellular trafficking (Hailey et al., 2017). G418-TR in saline (2 mg/kg), TR-hydrazide (2 mg/kg; Invitrogen, Waltham, MA, USA), or sterilized saline were injected s.c. into P5 neonatal mice and P25-30 adult mice. Neonatal mice were returned to cages with dams and euthanized 30 min to 72 h following injection. At time of euthanasia, whole blood was collected, and otic capsules were isolated in cold PBS. Otic capsules were fixed in 4% paraformaldehyde in PBS for 1 h at room temperature in the dark. Tissue was washed 3x with PBS, and the organ of Corti was dissected in PBS at room temperature. Basal, middle, and apical turns were isolated and transferred to a 48-well plate. The tissues were permeabilized and immunolabeled with anti-myosin VIIa antibodies (1:1000, Proteus Biosciences, Ramona, CA, USA) in 0.1% Triton X-100, 10% normal horse serum, and PBS overnight at 4 °C. After washing with PBS 3x, tissues were incubated with goat anti-rabbit Alexa 488 secondary antibody (1:750, Invitrogen, Waltham, MA, USA) and DAPI (1:2500, Thermo Scientific, Waltham, MA, USA) for 90 min at room temperature. Tissues were washed with PBS 3x and mounted in Fluoromount-G (Southern Biotech, Birmingham, AL, USA).

A similar procedure was carried out in adult animals. P25-30 mice were injected subcutaneously G418-TR in saline (2 mg/kg), TR-hydrazide (2 mg/kg), or sterilized saline. Animals were euthanized 6 h after injection and whole blood was collected. Otic capsules were removed and immediately transferred to cold 4% PFA. An insulin syringe with 27G needle was used to carefully make an apical cochleostomy, the round window was then pierced with the needle, and cold PFA was slowly infused into the cochlea. Otic capsules were then fixed at room temperature for 90 min. Tissues were washed with PBS 3x and then transferred to 500 mM EDTA for overnight decalcification at 4 °C. Tissue was washed with PBS, and the organ of Corti was dissected in PBS at room temperature. Immunolabeling and staining was the performed as above for the neonates.

### 2.4 ORC-13661 uptake inhibition

ORC-13661 was obtained from Oricula therapeutics. Compound was dissolved in sterile saline immediately prior to use. Animals were injected i.p. with ORC-13661 at 10-100mg/kg two hours prior to receiving aminoglycoside treatment. After two hours, animals were dosed with G418-TR s.c. at 10 mg/kg. Animals were returned to their cages, and then sacrificed at 3 and 6 hours after receiving G418-TR. For booster experiments animals received a booster dose of ORC-13661 3 hours following G418-TR administration. Animals and tissues were processed as described above.

### 2.5 Cryostat sectioning

Cochlea, kidneys, and livers were collected for cryostat sectioning. Animals were treated as indicated above, and after sacrifice the otic capsule was dissected in cold PBS and fixed in 4% PFA for 1 hour. Kidneys were also removed from animals at the time of sacrifice and immerse in 10% formalin. After washing in PBS, samples were sequentially dehydrated and equilibrated in 30% sucrose, 1:1 sucrose to OCT compound, and OCT compound overnight. Embedded tissue was frozen at -18°C and sectioned at a thickness of 12 µm. Slides were prepared with sectioned tissue. Cochlea, kidneys, and livers were labelled with phalloidin-Alexa 488 (1:200, Invitrogen, Waltham, MA, USA) for 2 hours at room temperature and DAPI (1:100) for 45 minutes at room temperature. Cochlea sections were also immunolabeled with rabbit anti-β3 tubulin (1:1000; clone TUJ1, Biolegend, San Diego, CA, USA) overnight at 4 °C followed by goat anti-rabbit Alexa-488 (1:200, Invitrogen, Waltham, MA, USA) for 2 hours at room temperature and DAPI (1:100) for 45 minutes at room temperature. All dilutions were in 5% normal goat serum and 0.1% Triton X-100 in PBS.

### 2.6 Confocal imaging

Slides were imaged with a Zeiss LSM 880 microscope (Zeiss, Oberkochen, Germany). For whole mount specimens, a 40x water objective was used to obtain 8-bit 1024 × 1024 pixel images of the mid-basal, mid-middle, and mid-apical turns. Confocal images were obtained using identical imaging parameters to compare fluorescence between images. For cryostat sectioned tissues, a 40x water objective was used to obtain 8-bit 1024 × 1024 pixel images of the organ of Corti, stria vascularis, spiral ganglia, and renal cortex. Identical imaging parameters were used for cochlea tissues, but laser intensity was decreased to avoid oversaturation when imaging renal cortex.

### 2.7 Fluorescence quantification

For whole mount fluorescence quantification, ImageJ software was used for image processing and fluorescence quantification. For neonatal hair cells, a single 256 × 165 pixel region of interest (ROI) was selected from the stack in the z-plane that captured all three rows of outer hair cells between the nucleus and cuticular plate. A similar ROI measuring 256 × 55 pixels was selected for the inner hair cells, again at a z-plane between the nucleus and cuticular plate. IHC and OHC ROIs were processed with a custom macro based on previously reported image analysis techniques (Makabe et al., 2020). Otsu thresholding was applied to the green channel to obtain a hair cell specific mask. This mask was then applied to the red channel image, and hair cell ROI mean fluorescence was calculated.

For cryostat sectioned tissue Phalloidin was used as a marker for hair cells, pillar cells, and stria vascularis tissue. OHC fluorescence was quantified by manually drawing an ROI for the region between the nucleus and the apical stereocilia. The ROI was then applied to the red channel, and mean fluorescence was quantified. Similarly, a manually segmented ROI for the pillar cells was used to quantify PC mean fluorescence. The stria vascularis ROI was also manually generated, and care was taken not to include acellular regions such as capillary lumens. For spiral ganglia neuron uptake quantification of tissue stained with Tuj1, a custom macro was generated that applied Otsu thresholding to the green channel to generate a mask which was then applied to the red channel to quantify mean fluorescence.

### 2.8 Statistics

Statistical analyses were done using GraphPad Prism 9.1.2. All statistics were performed comparing values from individual animals. When both cochlea were examined a mean value for the animal was calculated and used for analysis. Specific analyses are indicated in the figure legends. ANOVA was used to evaluate dose responses. Two-way ANOVA was used when multiple variables were analyzed for effect. When tissues from the same animal was compared, sample matching and a mixed effects model was used to account for any missing values. P-values less than 0.05 were considered significant.

### 2.9 Study approval

All experiments were approved by the University of Washington Institution Animal Care and Use Committee.

## 3 Results

### 3.1 Dose-dependent uptake of systemically administered G418-TR in neonatal mouse hair cells

To assess whether mammalian hair cells take up systemically injected geneticin (G418) tagged with Texas Red (TR), neonatal P5 C57/BL6 mice were subcutaneously (SC) injected with G418-TR (2.5-10 mg/kg), Texas Red-hydrazide (10 mg/kg) as a non-ototoxic control, or saline and sacrificed 6 hours after injection. The selected doses of G418-TR were not toxic to neonatal mice and hair cell damage was not evident in cochlear whole mount preparations. G418-TR, like gentamicin-TR (GTTR) (Dai et al., 2006; Makabe et al., 2020; Wang and Steyger, 2009), was detectable within neonatal hair cells 6 hours following SC injection with confocal microscopy of whole mount preparations (Figure 1A). Confocal microscopy demonstrates a dose-dependent increase in hair cell cytoplasmic accumulation of G418-TR. Control animals injected with 10 mg/kg unconjugated TR hydrazide demonstrate no significant hair cell fluorescence (Supplemental Figure 1A).

**Figure 1.**
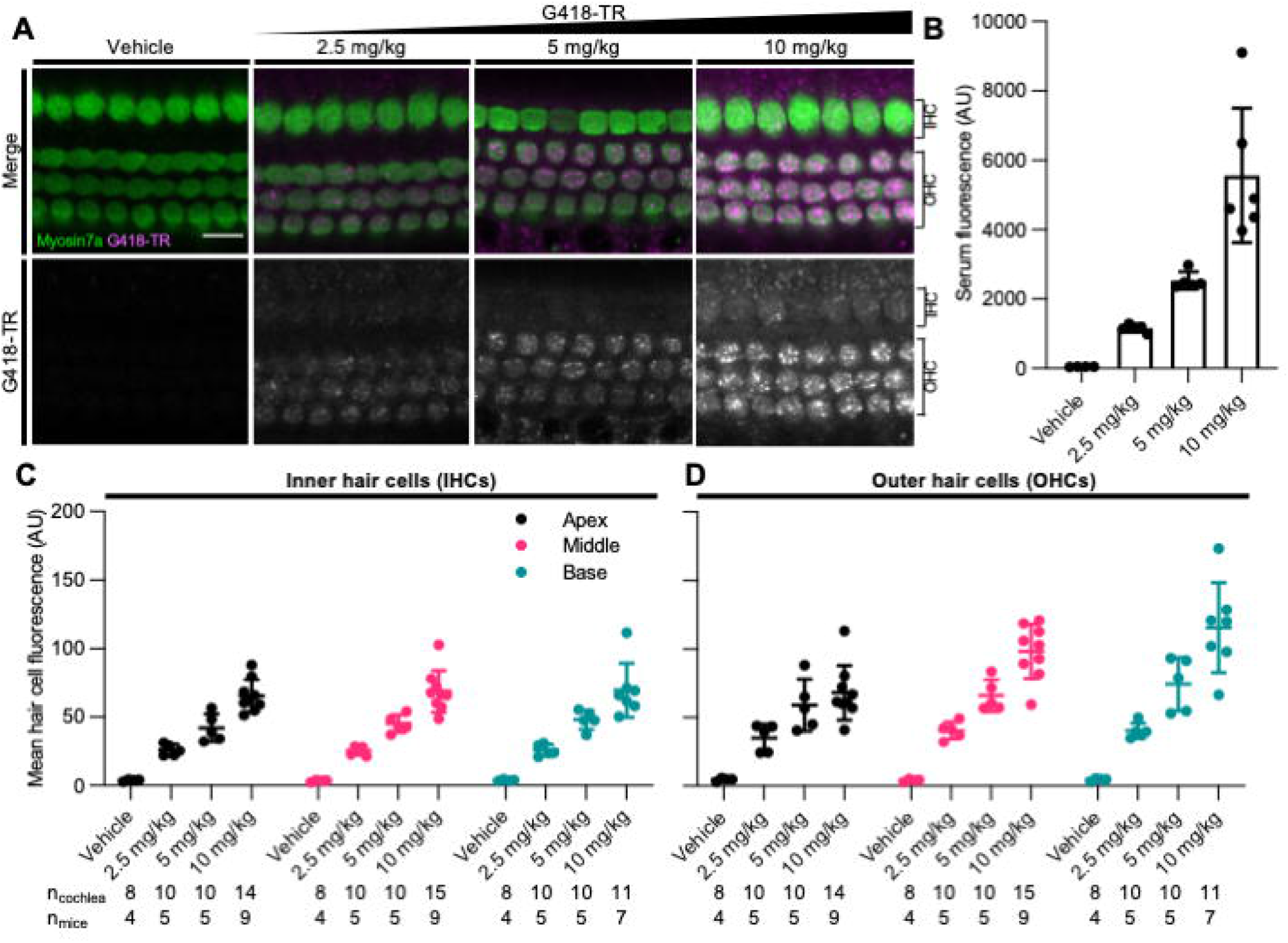
G418-TR is taken up by neonatal mammalian hair cells following systemic administration. (**A**) At 6 hours after systemic injection with G418-TR, neonatal mice (P5) were sacrificed, cochlear sensory epithelia was dissected and fixed, and tissue was immunolabeled for myosin VIIa. Representative single z-plane images captured from the middle turn of the cochlea in the region between the cuticular plate and nucleus demonstrate dose-dependent uptake of G418-TR in OHCs. Scale bar = 10 µm. (**B**) 6 hours after systemic administration of G418-TR, blood was collected from neonatal mice, serum was extracted, and fluorescence was quantified with a fluorescence plate reader. Increasing doses of G418-TR results in a dose-dependent increase in serum fluorescence (One-way ANOVA; F(_3,16_)=25.02, P<0.0001). (**C**) Mean G418-TR fluorescence intensity of IHCs by tonotopic region 6 hours after systemic treatment. IHCs demonstrate similar dose-dependent uptake of G418-TR in base, middle, and apex. (**D**) Mean G418-TR fluorescence intensity of OHCs by tonotopic region 6 hours after systemic treatment. OHCs demonstrate significant dose-dependent and tonotopic G418-TR uptake (Mixed effects analysis; F_(6,36)_=5.127, P=0.0007). n_cochlea_ and n_mice_ represent the number of cochlea and mice examined, respectively. When two cochlea from the same animal were analyzed, the animal mean was calculated and used for all statistics and graphs. Error bars are standard deviations. AU = arbitrary unit.

In order to quantify the dose-dependent uptake of G418-TR, we measured TR fluorescence present in mouse sera and cochlear hair cells. At time of euthanasia, blood was collected, serum was separated, and fluorescence was analyzed. A significant dose-dependent increase in serum fluorescence was observed in all treated animals (Figure 1B) after six hours of treatment. Control animals treated with TR-hydrazide demonstrated serum fluorescence that was not significantly different when compared to G418-TR treated animals (Supplemental figure 1B).

Quantification of inner hair cell (IHC) and outer hair cell (OHC) mean fluorescence was performed utilizing myosin VIIa immunolabeling as a hair cell-specific mask. Hair cells were analyzed from the apical, middle, and basal turns of each cochlea to assess for tonotopic differences in uptake. Following systemic administration of G418-TR in neonatal mice, both IHCs and OHCs demonstrated dose dependent increases in hair cell fluorescence (Figures 1C & 1D). When evaluating IHCs, all regions of the cochlea demonstrate a dose-dependent increase in uptake. Mixed-effect analysis with matching across tonotopic regions demonstrates a significant dose-effect (P<0.0001), but no significant variation in IHC uptake across tonotopic regions (P = 0.1090). OHCs demonstrated dose-dependent uptake across base, middle, and apex. Mixed-effect analysis with matching across tonotopic regions demonstrates significant interaction between dose and tonotopic region (P<0.001). Hair cells from animals treated with TR-hydrazide showed minimal background fluorescence that was not significantly different when compared to vehicle treated animals (supplemental figure 1B).

### 3.2 Dose-dependent uptake of systemically administered G418-TR in mature mouse hair cells is similar to observed neonatal hair cell uptake

Rodent cochlear hair cells are immature at birth and only begin to acquire the morphological and biophysical specializations characterizing mature hair cells during postnatal development (He et al., 1994; Marcotti and Kros, 1999; Abe et al., 2007; Jeng et al., 2020). Hearing onset in mice is generally thought to begin at approximately P12 and roughly corresponds with maturation of specializations related to hair cell type and location as well as other biophysical properties such as the endocochlear potential. In mice, the cochlea appears fully mature between P25-30 (Rubel, 1978; Kros et al., 1998; Marcotti et al., 2003; Rubel and Fay, 2012). To evaluate whether G418-TR uptake in mature hair cells was similar to that observed in neonatal mouse hair cells, juvenile C57/BL6 mice (25-30 days old) were subcutaneously (SC) injected with G418-TR (5-20 mg/kg), Texas Red-hydrazide (20 mg/kg), or saline and sacrificed 6 hours after injection.

Similar to neonatal hair cells, we observed that juvenile mouse hair cells readily take up systemically injected G418-TR (Figure 2a). Quantification of fluorescence from serum samples showed a significant dose-dependent increase in serum fluorescence (P<0.0001) (Figure 2B.). Both juvenile IHCs and OHCs demonstrated a dose-dependent increase in fluorescence across all tonotopic regions examined. Mixed-effect analysis with matching across tonotopic regions demonstrated significant dose-effects for IHCs or OHCs (P<0.001 and P<0.0001, respectively) but no significant variation from by tonotopic region for IHCs or OHCs (P=0.5924 and P=9312). Animals treated with TR-hydrazide demonstrated minimal fluorescence over background, which was similar to that observed in vehicle treated animals (Supplemental Figure 1C and 1D).

**Figure 2.**
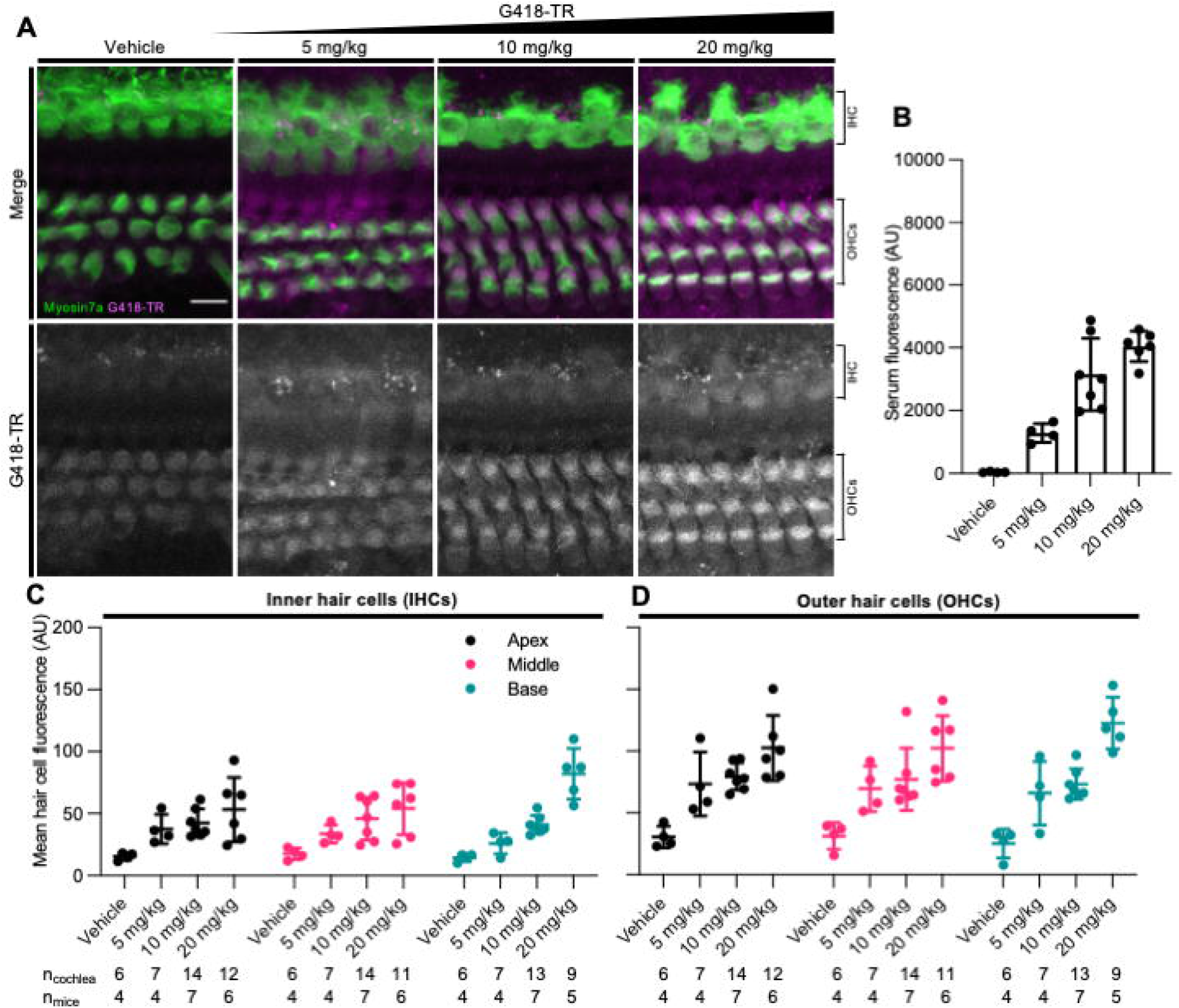
G418-TR is taken up by mature mammalian hair cells following systemic administration. (**A**) At 6 hours after systemic injection with G418-TR, juvenile mice (P25-30) were sacrificed, cochlear sensory epithelia was dissected and fixed, and tissue was immunolabeled for myosin VIIa. Representative maximum projection images captured from the middle turn of the cochlea demonstrate dose-dependent uptake of G418-TR in inner and outer hair cells. Scale bar = 10 µm. (**B**) 6 hours after systemic administration of G418-TR, blood was collected from juvenile mice, serum was extracted, and fluorescence was quantified with a fluorescence plate reader. Increasing doses of G418-TR results in a dose-dependent increase in serum fluorescence (One-way ANOVA; F_(3,17)_=28.28, P<0.0001). (**C**) Mean G418-TR fluorescence intensity of IHCs by tonotopic region 6 hours after systemic treatment. IHCs demonstrate low uptake at most doses of G418-TR with no significant tonotopic variation. (**D**) Mean G418-TR fluorescence intensity of OHCs by tonotopic region 6 hours after systemic treatment. OHCs demonstrate dose-dependent uptake but not tonotopic variation in G418-TR uptake. n_cochlea_ and n_mice_ represent the number of cochlea and mice examined, respectively. When two cochlea from the same animal were analyzed, the animal mean was calculated and used for all statistics and graphs. Error bars are standard deviations. AU = arbitrary unit.

### 3.3 Characteristics of G418-TR hair cell accumulation

Aminoglycoside-induced hair cell toxicity typically initially occurs in basal OHCs, and extends apically and to IHCs with increasing cumulative dosing (Forge and Schacht, 2000). We sought to determine whether this pattern of differential toxicity correlated with levels of G418-TR uptake. In P5 neonatal mice treated with 10 mg/kg G418-TR for six hours, we observed dramatic tonotopic variation in uptake in OHCs, but not IHCs (Figure 3A). Basal turn OHCs demonstrated nearly twice the mean fluorescence compared to apical OHCs (115.8 AU and 68.2 AU, respectively). However, IHCs from the basal, middle, and apical turns from the same animals showed similar levels of fluorescence (69.7 AU, 68.8 AU, and 65.5 AU, respectively). In older mice with mature hearing (P25-30) treated with 20 mg/kg G418-TR for six hours, we observed a much small but significant increase in IHC and OHC fluorescence in the basal turn only (Figure 3C).

**Figure 3.**
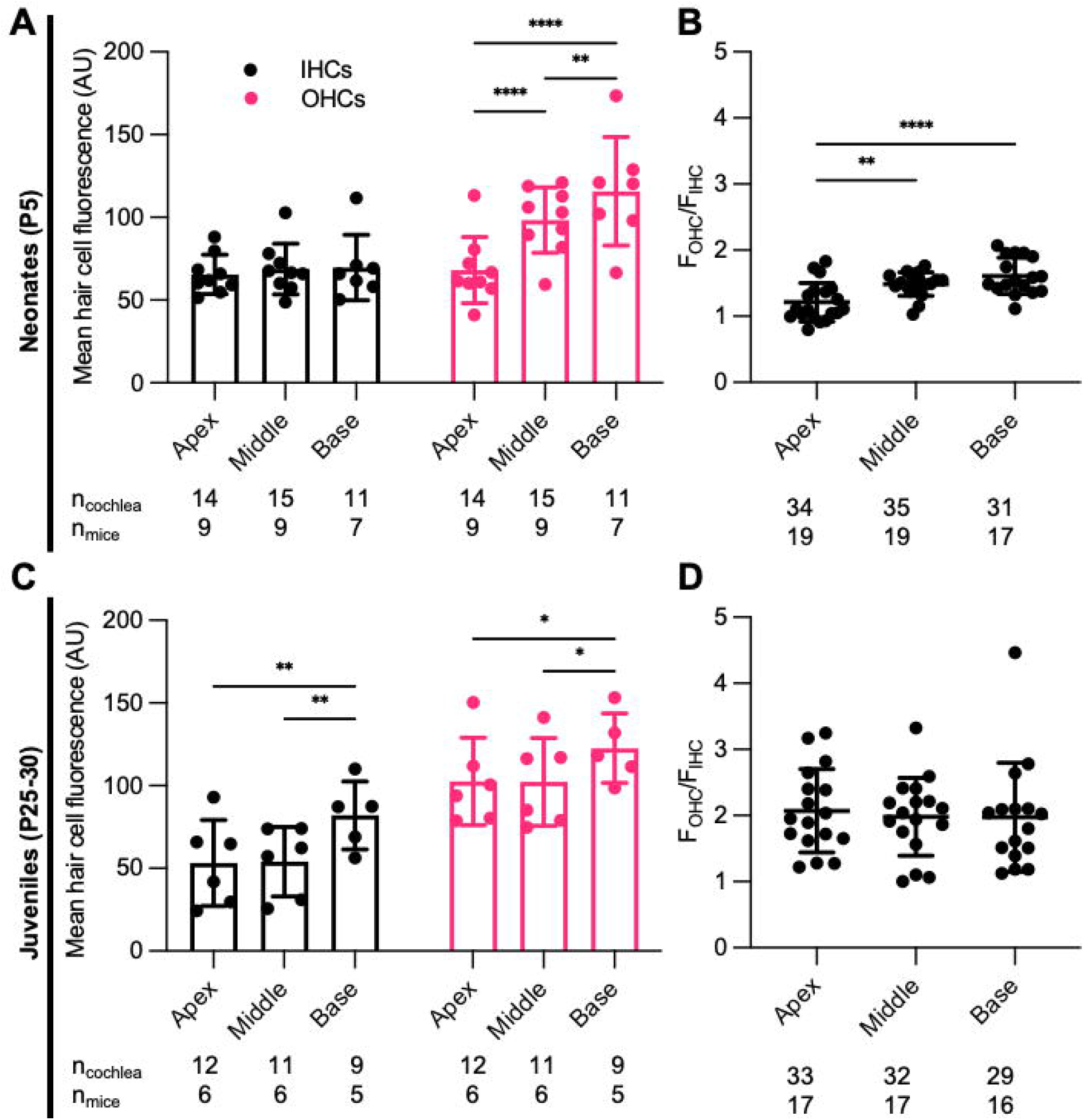
Characteristics of G418-TR uptake in neonatal and juvenile hair cells. (**A**) 6 hours after systemic injection with G418-TR, tonotopic variation in uptake is observed in neonatal hair cells. Neonatal (P5) mice treated with 10 mg/kg G418-TR show no significant tonotopic variation in IHC uptake (Mixed-effects Analysis; F(_2,14_)=2.731, P=0.0996) but significant variation in OHC uptake (Mixed-effects Analysis; F(_2,14_)=17.05, P=0.0002). (**B**) Pooled analysis of the ratio of OHC to IHC uptake (F_OHC_/F_IHC_) was calculated for all neonatal animals treated with G418-TR (2.5, 5, and 10 mg/kg) and sorted by tonotopic region. In neonatal hair cells, there is significant variation in F_OHC_/F_IHC_ by region (Mixed-effects Analysis; F(_2,52_)=11.81, P<0.0001). (**C**) Juvenile mice (P25-30) treated with 20 mg/kg G418-TR show tonotopic variation in uptake in both IHCs (Mixed-effects Analysis; F(_2,09_)=10.77, P=0.0041) and OHCs (Mixed-effects Analysis; F(_2,09_)=5.394, P=0.0289). (**D**) Pooled analysis for all juvenile animals treated with G418-TR (5, 10, and 20 mg/kg) demonstrates no significant tonotopic variation in F_OHC_/F_IHC_ (Mixed-effects Analysis; F(_2,31_)=0.1442, P=0.8662). n_cochlea_ and n_mice_ represent the number of cochlea and mice examined, respectively. When two cochlea from the same animal were analyzed, the animal mean was calculated and used for all statistics and graphs. Error bars are standard deviations. AU = arbitrary unit.*P<0.05, **P<0.01, ***P<0.001, ****P<0.0001; P values generated from multiple comparisons post-hoc test (Tukey’s multiple comparison test).

In each cochlea we directly compared the ratio of OHC fluorescence to IHC fluorescence (F_OHC_/F_IHC_) in each tonotopic region. Apical, middle, and basal hair cells demonstrated significantly higher fluorescence in OHCs compared to IHCs (Figure 3B & 3D). When we analyzed data from all neonates treated with G418-TR (2.5-10 mg/kg) for six hours, the ratio between OHC and IHC uptake increased from a mean of 1.21 at the apex to 1.61 at base. When the same analysis was applied to specimens collected from older animals (P25-30), mean F_OHC_/F_IHC_ were 2.18, 2.11, and 2.02 in the apex, middle and base, respectively. Note as well that only 5 of 55 neonatal cochlea samples and 0 of the 50 juvenile cochlea samples examined in this way had F_OHC_/F_IHC_ ratio < 1.0.

### 3.4 Neonatal hair cells accumulate and retain G418-TR

We also studied the relationship between G418-TR serum kinetics and hair cell uptake. We treated neonatal P5 mice with 10 mg/kg G418-TR and sacrificed cohorts of animals 30 minutes – 72 hours after injection. At the time of sacrifice, whole blood was collected from animals and serum was isolated for fluorescence quantification. Serum fluorescence was converted to concentration using a calibration curve of G418-TR diluted in fetal bovine serum. Serum kinetics for G418-TR demonstrate a peak at 3 hours post-injection with a t_1/2_ of 2.5 hours, resembling the parameters previously reported for gentamicin-TR (Wang and Steyger, 2009) (Figure 4B).

**Figure 4.**
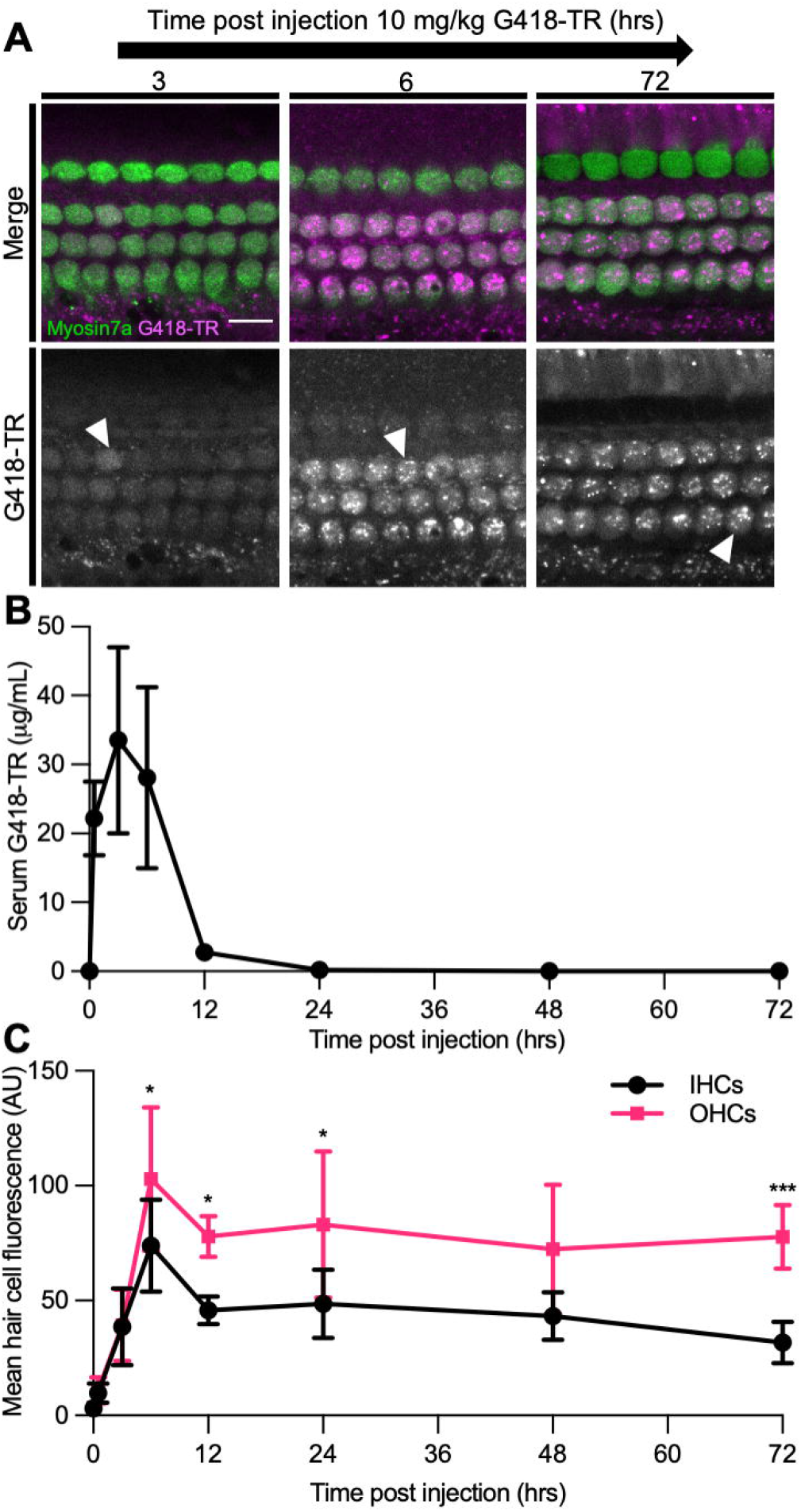
Hair cells accumulate and retain G418-TR after systemic injection. (**A**) 30 minutes-72 hours after systemic injection with 10 mg/kg G418-TR, neonatal mice (P5) were sacrificed, cochlear sensory epithelia was dissected and fixed, and tissue was immunolabeled for myosin VIIa. Representative images captured from the middle turn of the cochlea demonstrate uptake and retention of G418-TR. Images shown are single z-plane images capturing the region between the nucleus and cuticular plate. Arrowhead highlights transition from diffuse G418-TR distribution to accumulation in puncta with time. Scale bar = 10 µm. (**B**) Serum kinetics of G418-TR after systemic injection. After systemic injection with 10 mg/kg G418-TR, neonatal mice were sacrificed at different time points, serum was collected, and fluorescence was quantified. Fluorescence was converted to concentration in µg/mL utilizing a calibration curve of G418-TR diluted in fetal bovine serum. Quantification shows that serum levels of G418-TR peak at 3hrs and are undetectable 24 hours post-injection. (**C**) Mean hair cell fluorescence from the middle turn of the cochlea was calculated over 72 hours. Both IHCs and OHCs show uptake and retention. Animals examined (t [hours], n) = (0,4), (0.5,4), (3,5), (6,6), (12,5), (24,5), (48,5), (72,5). Error bars are standard deviations. AU = arbitrary unit. Significant differences between IHC and OHCs denoted with: *P<0.05, **P<0.01, ***P<0.001. P values calculated from multiple comparisons post-hoc test (Šídák’s multiple comparison test).

We also evaluated the kinetics of hair cell uptake and accumulation of G418-TR in hair cells from the same animals over 72 hours. After subcutaneous injection with 10 mg/kg G418-TR, both OHCs and IHCs accumulate G418-TR over 6 hours, and then retain that fluorescence for at least 72 hours post-injection (Figure 4A). Quantification of middle turn hair cell fluorescence demonstrates that IHC and OHC fluorescence peaks at 6 hours, and then plateaus with no significant decrease from 12 – 72 hours post-injection (Figure 4C). IHC and OHC fluorescence is similar for the first 3 hours, but after six hours the difference between IHC and OHC mean fluorescence increases and remains significantly different. Statistical analysis demonstrates significant interaction between time since injection and hair cell population (two-way ANOVA, F_(7,62)_=2.440; P<0.05). Post-hoc testing for multiple comparisons demonstrates significant differences between IHC and OHC values at 6, 12, 24, and 72 hours post injection (P<0.05, P<0.05, P<0.05, and P<0.001, respectively).

### 3.5 G418-TR accumulation in the kidney

Aminoglycoside nephrotoxicity is attributed to cytotoxic affects within the tubular epithelium (Lopez-Novoa et al., 2011). Within the nephron, the proximal convoluted tubule (PCT) cells, but not the glomerulus or distal convoluted tubule (DCT) cells, show unique susceptibility to aminoglycosides (Mingeot-Leclercq and Tulkens, 1999). Previous work using GTTR demonstrated that PCT cells rapidly take up aminoglycoside in vivo (Dunn et al., 2002; Dai et al., 2006). Kidneys were harvested from animals treated with 10 mg/kg G418-TR in order to evaluate renal aminoglycoside uptake and retention over 72 hours (Figure 5). Similar to findings for hair cells, PCT epithelial cells rapidly accumulate G418-TR over 24 hours and retain that fluorescence for up to 72 hours. DCT cells and glomeruli show less accumulation of G418-TR, but it should still be noted that these cell populations demonstrate measurable G418-TR accumulation and retention. Additionally, it is worth noting that the intensity of the G418-TR signal was significantly brighter within the kidney compared to the cochlea, and the excitation laser intensity needed to be decreased in order to prevent over saturation of the detectors used. Because aminoglycosides are renally excreted, the nephrons of the kidney are exposed to significantly more aminoglycoside than the cochlea, accounting for this notable difference in overall fluorescence levels.

**Figure 5.**
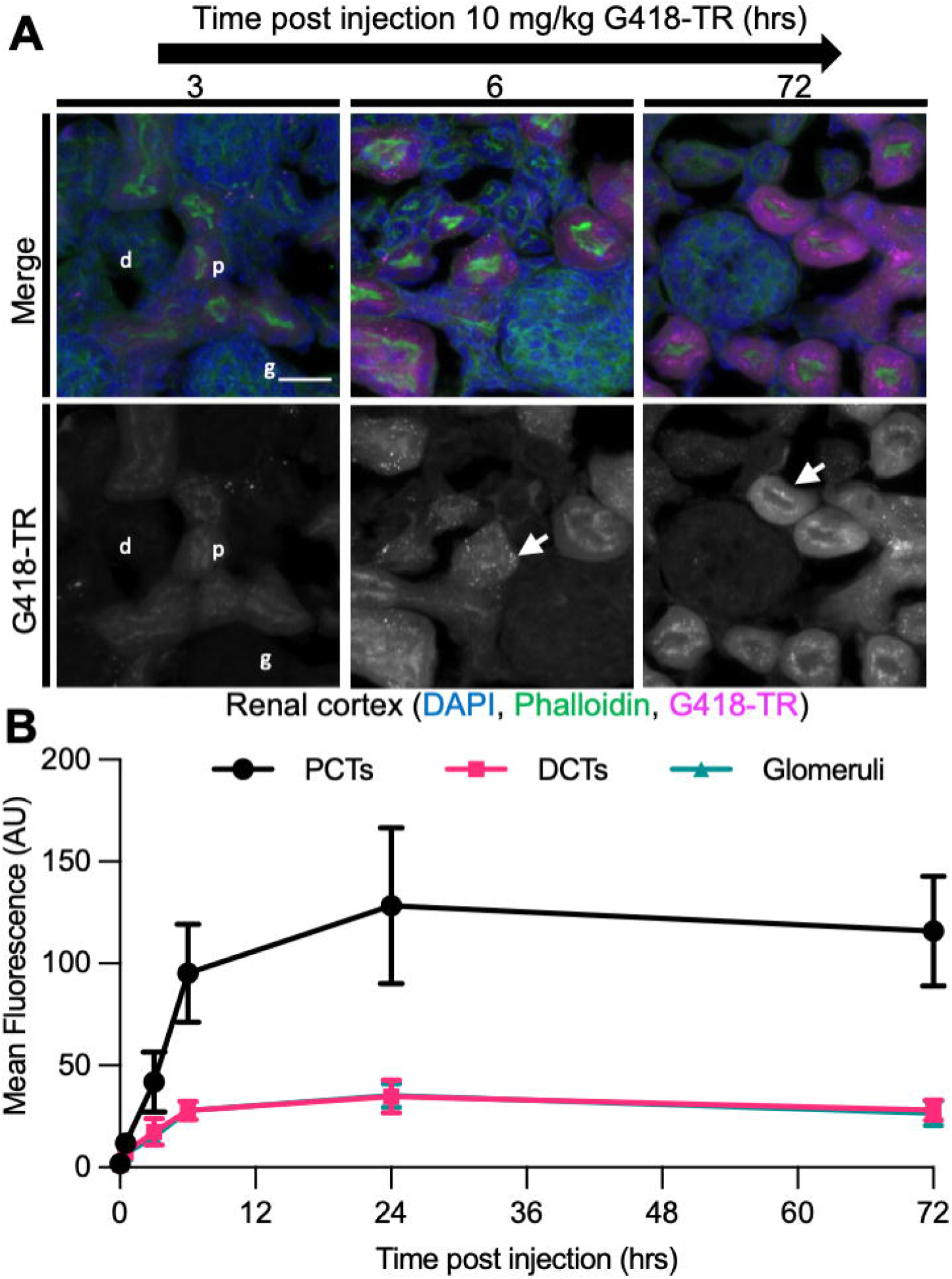
Selective aminoglycoside retention in proximal convoluted tubule cells of the kidney. **(A)**0 minutes-72 hours after systemic injection with G418-TR, neonatal mice (P5) were sacrificed, and kidneys were removed. Tissues were fixed, embedded and cryostat sectioned at a thickness of 12 µm. Tissue sections were stained with phalloidin and DAPI. Representative maximum projection images from the renal cortex show proximal convoluted tubule (p) uptake and retention, but minimal uptake in the distal convoluted tubule (d) and glomeruli (g). Like hair cells, PCT cells also show apical puncta (arrow) accumulation of G418-TR. Scale bar = 20 µm. (**B**) Mean fluorescence from proximal convoluted tubules (PCTs), distal convoluted tubules (DCTs), and glomeruli was calculated over 72 hours. PCTs demonstrate G418-TR uptake and retention like that observed in cochlear hair cells. Animals examined n = 4 for each time point. Error bars are standard deviations. AU = arbitrary unit.

Aminoglycoside hepatotoxicity is rare (Aminoglycosides, 2012). Whether this is due to dose limiting toxicity in other organs (e.g. nephrotoxicity) or an intrinsic property of the liver that renders it less susceptible to aminoglycoside toxicity is unknown. When livers from animals treated with G418-TR were examined, we observed low levels of fluorescence analogous to that observed in the DCT and glomeruli of the kidney (Supplemental Figure 2). This complements previous analyses that noted low-level aminoglycoside uptake in the liver of animals treated with radioisotope labeled aminoglycosides (Huy et al., 1986).

### 3.6 G418-TR accumulation in non-sensory cochlear tissues

In addition to hair cells, systemically administered aminoglycosides have been shown to accumulate in the non-sensory tissues of the inner ear (Bareggi et al., 1986; Imamura and Adams, 2003; Kitahara et al., 2005; Wang and Steyger, 2009). In neonatal mice treated with a single dose of 10 mg/kg G418-TR, we observed fluorescence in cryostat sectioned cochlear tissues in the organ of Corti, stria vascularis, and spiral ganglia neurons (Supplemental Figure 3).

The organ of Corti includes both the mechanosensory hair cells as well as numerous supporting cell populations and neuronal tissues. Analysis of OHCs in cross section supports the findings reported above for whole mount images; OHCs accumulate G418-TR in a diffuse pattern over 6 hours, and at later time points G418-TR fluorescence is sequestered in apical regions (Supplemental Figure 3A). In neonatal mice, we observed a transient increase in stria vascularis fluorescence following systemic administration (Supplemental Figure 3B). Aminoglycosides have also been localized to spiral ganglia neurons following systemic injection (Imamura and Adams, 2003; Kitahara et al., 2005). Unlike the vascular stria vascularis, spiral ganglia neurons demonstrated consistent low-level fluorescence (Supplemental Figure 3C).

To compare the relative uptake of different tissues within the cochlea, we quantified mean tissue fluorescence at 0, 6, 24, and 48 hours in each region (Figure 6). OHCs show a similar pattern as shown in whole mount imaging and quantification (Figure 4), with a peak at 6 hours followed by retention of AG at later time points. Stria vascularis fluorescence demonstrated a peak in fluorescence at 6 hours post injection followed by a decrease to near background levels by 48 hours. Pillar cells, located between IHCs and OHCs demonstrate relatively low-level uptake and retention of G418-TR, while spiral ganglion neurons demonstrated minimal uptake at all time points.

**Figure 6.**
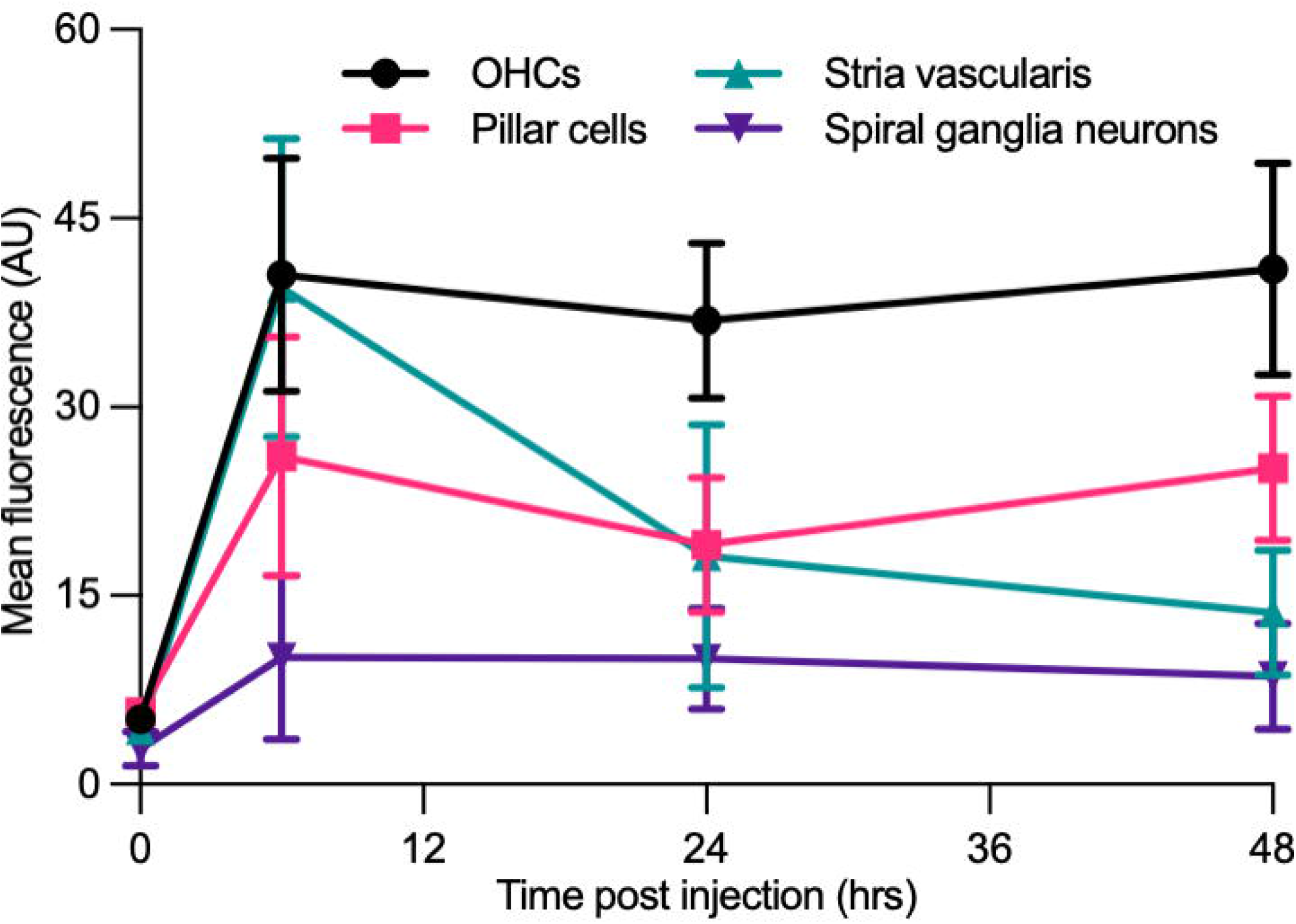
Differential aminoglycoside uptake and retention within the cochlea. (**A**) Cochlea from neonatal mice treated with G418-TR were fixed, cryostat sectioned, and signed in order to analyze the organ of Corti, stria vascularis, and spiral ganglia neurons. In order to compare relative G418-TR uptake between specific tissue types, fluorescence was quantified for outer hair cells (OHCs), pillar cells, stria vascularis, and spiral ganglia neurons. Note that while OHCs take up and retain G418-TR fluorescence, stria vascularis tissue has an early peak but then declines while SGNs show consistent low level uptake. PCs display an intermediate level of uptake with some retention. Error bars are standard deviations. Animal numbers (time [hours], n) = (0,3), (6,4), (24,4), and (48,4).

### 3.7 ORC-13661 blocks the uptake of G418-TR in vivo in mammalian hair cells

ORC-13661 is a novel therapeutic that protects hair cells from aminoglycoside toxicity in fish and mammals (Chowdhury et al., 2018; Kitcher et al., 2019). Mechanistic studies in zebrafish and mouse cochlear cultures demonstrate that ORC-13661 reversibly blocks mechanoelectrical transduction (MET) channel currents and uptake of fluorescently labeled aminoglycosides (Kitcher et al., 2019). To test whether ORC-13661 blocks uptake of aminoglycosides in vivo in mice, neonatal mice were pre-treated with saline or ORC-13661 (10-100 mg/kg; i.p.). Two hours after this treatment mice were injected with G418-TR (10 mg/kg; s.c.). Animals were sacrificed at either 3 or 6 hours after exposure to G418-TR to test both 3 hour (“short”) and 6 hour (“long”) G418-TR exposure paradigms (Supplemental Figure 4, A & B).

Analysis of the mid-basal turn hair cells demonstrated inhibition of uptake by ORC-13661 in both treatment schemes (Figure 7, A & B). When animals are pre-treated with vehicle and injected with G418-TR, robust hair cell uptake is observed in both treatment schemes. When pretreated with increasing doses of ORC-13661 followed by injection with G418-TR, the short G418-TR exposure scheme demonstrates significant uptake inhibition at all doses of ORC-13661. Interestingly, the long G418-TR exposure scheme showed uptake inhibition at only the 100 mg/kg ORC-13661 dose.

**Figure 7.**
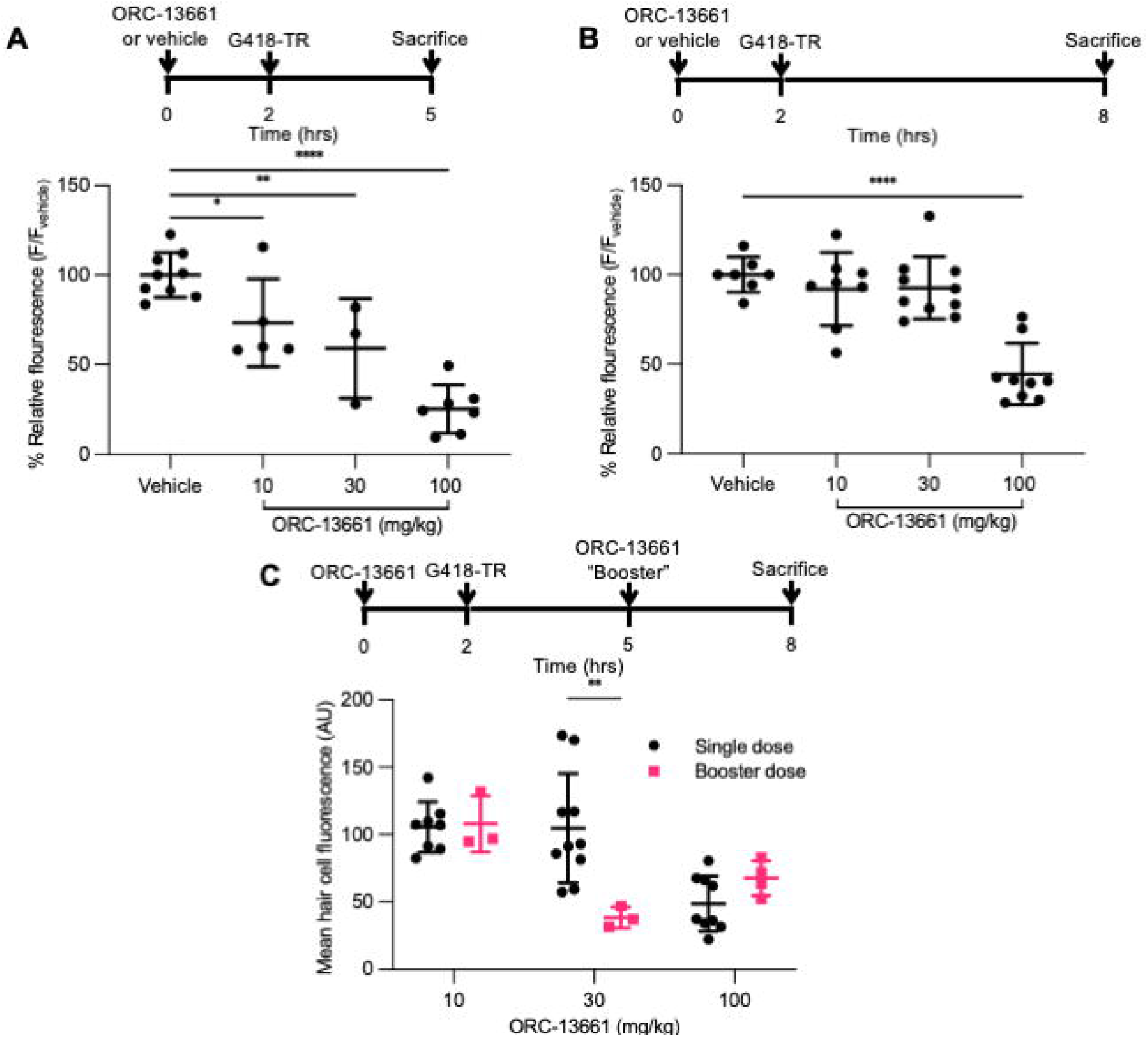
ORC-13661 blocks G418-TR accumulation in mammalian hair cells. (**A**) Short (3 hour) G418-TR exposure paradigm. Neonatal mice were pretreated with ORC-13661 or vehicle for 2 hours, injected with 10 mg/kg G418-TR, and sacrificed 3 hours after aminoglycoside injection. Basal turn OHC mean fluorescence for animals treated with either vehicle or ORC-13661 followed by 10mg/kg G418-TR. OHC mean fluorescence was quantified and normalized to animals pre-treated with vehicle within each experimental replicate. (One-way ANOVA; F(_3,20_)=23.55, P<0.0001). (**B**) Long (6 hour) G418-TR exposure paradigm. Neonatal mice were pretreated with ORC-13661 or vehicle for 2 hours, injected with 10 mg/kg G418-TR, and sacrificed 6 hours after aminoglycoside injection. (One-way ANOVA; F(_3,30_)=19.62, P<0.0001). (**C**) Booster dosing strategy with a single pretreatment injection of ORC-13661 followed by a second booster dose 3 hours after G418-TR injection. Basal turn OHC mean fluorescence for animals treated with a pretreatment dose ORC-13661 (10-100 mg/kg) followed by 10 mg/kg G418-TR with an additional booster of ORC-13661 (same as pretreatment dose) 3 hours after aminoglycoside administration. Note that data for single dose are non-normalized data from part B. Error bars are standard deviations. *P<0.05, **P<0.01, ***P<0.001, ****P<0.0001; P values generated from multiple comparisons post-hoc test (A&B: Dunnett’s multiple comparison test; C: Šídák’s multiple comparison test).

To compare results between experiments, hair cell fluorescence was normalized to intra-experimental vehicle treated animals. By normalizing within experimental groups, we were able to minimize the variability between experimental groups (as observed in neonates treated with 10 mg/kg G418-TR [Figure 1, C & D]) and discriminate smaller effects on hair cell uptake. We quantified relative fluorescence for mid-basal turn OHCs and observed a significant dose-dependent inhibition of uptake for increasing doses of ORC-13661 in the short exposure scheme with significant uptake inhibition compared to vehicle treated animals detected at all ORC-13661 dose levels (Figure 7A). For animals treated in the long exposure paradigm, significant uptake inhibition was only noted at the highest dose of ORC-13661 (Figure 7B). Kidneys from animals in the 100 mg/kg ORC-13661 dose group of the long exposure scheme demonstrated no change in PCT G418-TR uptake (Supplemental Figure 5). This supports the hypothesis that ORC-13661 protects hair cells through selective inhibition of MET-dependent aminoglycoside uptake, as opposed to endocytosis.

When comparing the two different treatment schemes with a two-way ANOVA, there is no significant difference in OHC uptake inhibition (F_(3,50)_=1.989, P=0.1276) however component analysis demonstrates a significant impact from dose (F_(3,50)_=39.75, P<0.0001) and treatment scheme (F_(1,50)_=13.45, P=0.0006). These findings suggest that inhibition of aminoglycoside uptake by ORC-13661 in neonatal mice is both dose- and time-dependent.

We hypothesized that the difference between the short and long exposure paradigms was due to different pharmacokinetics for G418-TR and ORC-13661. At 6 hours, we observed high levels or circulating G418-TR (Figure 4B). If the elimination kinetics of ORC-13661 in the neonatal mouse are rapid, this might explain the difference between the short and long G418-TR exposure paradigms. To test this hypothesis, we developed a “booster” treatment scheme. Neonatal mice were treated with a single pretreatment of ORC-13661 (10, 30, or 100 mg/kg), then injected with 10 mg/kg G418-TR, and given a booster dose of ORC-13661 (same dose as pretreatment) 3 hours after G418-TR (Figure 7C). When compared to animals that received a single pretreatment dose of ORC-13661, animals that received 30 mg/kg ORC-13661 pretreatment and a 30 mg/kg ORC-13661 booster showed significantly more uptake inhibition. The 10 mg/kg and 100 mg/kg ORC-1366 booster schemes showed no significant difference when compared to single doses, suggesting that the 10 mg/kg dose is not enough to generate uptake inhibition while the 100 mg/kg dose may be reaching a plateau and an additional booster does not increase uptake inhibition. These findings support our hypothesis and further demonstrating the translational utility of this in vivo system.

## 4 Discussion

Numerous drugs and therapeutic compounds are known to be toxic to the inner ear and adversely affect hearing and balance, however our understanding of the mechanisms driving ototoxicity remains incomplete. Challenges in studying ototoxicity have led to a paucity of protective therapeutics. The zebrafish lateral line and rodent explants have proven useful in screening and mechanistic studies, but translating these discoveries to adult mammals in vivo remains a difficult and costly endeavor. We present the development of low-cost and rapid in vivo mammalian model for studying ototoxic compounds and candidate protective therapeutics.

### 4.1 Biomarker

In mammals, diagnosis of ototoxicity is usually limited to functional evaluations such as behavioral paradigms, auditory brainstem response (ABR) thresholds, and otoacoustic emissions analysis or post-mortem histologic evaluation. These methods are expensive and time consuming, and they generally only detect ototoxic effects after irreversible damage has occurred. Some in vivo mammalian ototoxicity protocols also require cotreatment with loop diuretics, a class of therapeutics that are both independently ototoxic and show synergistic toxicity with other compounds (Rybak, 1993; Ding et al., 2016). While these protocols allow for rapid induction of in vivo ototoxicity, the introduction of a second therapeutic agent with independent effects on numerous organ systems confounds analysis and clinical relevance. Indeed, concomitant use of loop diuretics with aminoglycosides or cisplatin in humans is generally cautioned against in all but the most exceptional of circumstances (Huxel et al., 2021; Furosemide: Drug information - UpToDate). We hypothesized that by injecting neonatal mice with G418-TR, we could develop a rapid and sensitive technique for evaluating in vivo aminoglycoside toxicity. In this study, we demonstrate that systemic injection of G418-TR into neonatal mice results in quantifiable, dose-dependent hair cell uptake of the aminoglycoside. Further evaluation of tonotopic and cell specific uptake patterns demonstrates that basal regions of the cochlea take up more aminoglycoside compared to apical regions, and OHCs take up significantly more aminoglycoside than immediately adjacent IHCs. These findings are consistent with the hypothesis that differential aminoglycoside toxicity within the inner ear is due to differential exposure of the hair cells. Whether this increased exposure is due to characteristics of aminoglycoside trafficking within the cochlea or local hair cell specific properties such as MET channel permeability remains unknown.

Because mice are born with immature cochlear hair cells and functional hearing only develops around P12, we compared neonatal hair cell uptake with that of mature hair cells from juvenile animals P25-30. Slightly higher doses of G418-TR were required in juvenile animals to achieve a similar dose-dependent increase in hair cell aminoglycoside accumulation, however the apicobasal gradient and OHC predilection were again observed. The lower doses required in neonates is consistent with the previously described “sensitive period” during rodent development when neonates are more sensitive to aminoglycosides than older animals (Chen and Saunders, 1983; Nakai et al., 1983; Prieve and Yanz, 1984).

Additionally, it is worth noting that in both neonates and juveniles we observed no animal mortality and no significant hair cell death. This is due to the fact that our studies utilized sublethal dosing of G418-TR (maximum of 10 mg/kg in neonates and 20 mg/kg in juveniles) far lower than previously described aminoglycoside ototoxicity protocols that require multiple sequential doses of 0.5-1g/kg (Chowdhury et al., 2018; Wu et al., 2001; Horvath et al., 2019).

With similar hair cell uptake patterns observed in neonates and juveniles, albeit requiring slightly different dosing, we elected to pursue the remainder of our experiments in neonates. By using neonates we estimate we saved 80-90% on animal fees (housing only breeding animals), used 1/10th as much reagent (treating 2 g neonates vs. 20 g adults), and decreased our tissue processing time by about 50% (no decalcification required). The significant time and cost savings, coupled with faithful recapitulation of results observed in older animals, makes the neonate an attractive system for rapid and higher throughput translation of preclinical discoveries into mammalian systems designed to evaluate hearing loss and protection.

### 4.2 Characterization of in vivo mammalian hair cell uptake and retention of G418-TR

In mammalian cochlear explant models there is clear evidence that aminoglycosides act as permeant blockers of MET channels, rapidly gaining entry into hair cells and accumulating without means of egress (Marcotti et al., 2005; Alharazneh et al., 2011; Vu et al., 2013). Aminoglycoside uptake and retention has also been documented in vivo (Hiel et al., 1993; Imamura and Adams, 2003; Dai et al., 2006). We sought to quantify the in vivo mammalian kinetics of aminoglycoside uptake in hair cells. Neonatal hair cells demonstrate rapid uptake of G418-TR from 0-6 hours post-injection. Interestingly, we noted differences between IHC and OHC uptake and retention. Between 0-3 hours, IHCs and OHCs demonstrate nearly identical levels of uptake. From 3-6 hours post-injection, OHCs demonstrate a greater rate of uptake compared to IHCs which results in a significant difference in peak fluorescence between the two populations at 6 hours. After reaching peak hair cell fluorescence levels at 6 hours post-injections, OHCs demonstrate no significant loss by 72 hours while IHCs show a moderate reduction in fluorescence. When correlated with measured serum fluorescence, we noted that hair cell uptake corresponds to the peak in serum G418-TR levels; when circulating G418-TR is maximal, the hair cells readily accumulate aminoglycoside. Once serum levels of aminoglycoside begin to drop, OHCs retain aminoglycoside while IHCs demonstrate a slow loss of signal. The mechanism of aminoglycoside egress from IHCs is undetermined as these charged molecules are unlikely to cross membranes by passive diffusion.

Our studies are consistent with previous studies using autoradiography and immunolabeling that have established that hair cells rapidly take up aminoglycosides and retain them for extended durations (Portmann et al., 1974; de Groot et al., 1990; Imamura and Adams, 2003; Dai et al., 2006). Previous studies have noted that aminoglycoside uptake similarly shows a basal OHC predilection using qualitative and semi-quantitative methods of analysis (Hiel et al., 1993; Imamura and Adams, 2003; Dai et al., 2006). An almost universal finding is that outer hair cells are more susceptible to aminoglycoside ototoxicity than inner hair cells. This difference in ototoxicity remains poorly understood. It is sometimes attributed to the higher metabolic rate of outer hair cells (given their contribution to the active process of the cochlear amplifier), but there is also conflicting evidence for differential aminoglycoside uptake and accumulation in IHCs and OHCs (Lim, 1986; Hiel et al., 1993; Forge and Schacht, 2000; Imamura and Adams, 2003; Taylor et al., 2008; Wang and Steyger, 2009). Our findings support the hypothesis that differential aminoglycoside susceptibility between IHCs and OHCs is in part due to differential exposure. Explant studies have demonstrated differential rates of labeled aminoglycoside uptake in OHCs and IHCs (Alharazneh et al., 2011), but in vivo uptake of GTTR previously implied comparable IHC and OHC fluorescence at 3 hours post-injection (Wang and Steyger, 2009; Makabe et al., 2020). In agreement with these previous studies, our results demonstrate similar IHC and OHC uptake from 0-3 hours, but we find that a significant difference between OHCs and IHCs develops at 6 hours and persists out to 72 hours. We suspect that this difference is due in part to the difference in resting potentials between the hair cell populations, with a more negative OHC resting potential resulting in more favorable electrostatic uptake of positive aminoglycosides (Pickles, 1998). We also demonstrate that IHCs, unlike OHCs, show some level of aminoglycoside egress which could also be contributing to this differential susceptibility. It is worth noting that previous work has also suggested different isoforms of the MET channel could also contribute to differences between OHCs and IHCs (Beurg et al., 2006; Fettiplace, 2009), and certainly these differences could also impact aminoglycoside uptake. Quantification of aminoglycoside levels after an initial peak at 6 hours also allowed for evaluation of aminoglycoside retention in hair cells. This process is an ideal application of the neonate system, because hair cell addition and turnover in aquatic vertebrates complicates prolonged analysis and in adult mammals the cost of a cross sectional study with adequate numbers is prohibitive to most research programs. We also observe not only differential accumulation, but also clearance of aminoglycoside. Despite the fact that mechanosensory hair cells and renal proximal tubule cells are known to retain aminoglycoside(Imamura and Adams, 2003; Dai et al., 2006), aminoglycoside clearance at the cellular level has not been well studied. The biologic relevance of this observation is unclear, but certainly warrants additional investigation.

Histologic evaluation of the kidneys from G418-TR treated mice corroborates previous studies which demonstrated rapid and sustained aminoglycoside uptake by the PCT cells (Dunn et al., 2002; Dai et al., 2006). While quantification of fluorescence from the PCT reveals a similar pattern of uptake and retention compared to that observed in cochlear hair cells, qualitatively there are also similarities in the cellular distribution of G418-TR. Both PCTs and hair cells show a pattern of diffuse uptake at early time points, followed by what appears to sequestration in apical puncta. Despite these similarities, hair cells and PCTs are thought to uptake aminoglycosides via distinct pathways. Hair cells accumulate aminoglycosides primarily through MET channel transport, with a relatively minor contribution from endocytosis (Huth et al., 2011; Hailey et al., 2017; Makabe et al., 2020). In the nephron, mutation of the PCT endocytic receptor megalin blocks aminoglycoside uptake and protects against nephrotoxicity (Schmitz et al., 2002). Given the differences in uptake mechanisms, the similarities in aminoglycoside uptake and localization in hair cells and PCTs suggest there is convergence of similar mechanisms of intracellular aminoglycoside handling and sequestration. This hypothesis is supported by the observation that aminoglycosides accumulate in lysosomal compartments within both hair cells (Hailey et al., 2017; Hashino et al., 1997) and PCT cells (Silverblatt and Kuehn, 1979).

Evaluation of non-sensory tissues from the cochlea, including the supporting cells, spiral ganglia neurons, and the stria vascularis further demonstrates the unique pattern of uptake observed in hair cells. Cross sectional analysis of cryostat sections demonstrated once again that hair cells take up G418-TR over 6 hours and then retain the aminoglycoside load. Qualitatively, we again observe aminoglycoside entering the hair cells in a diffuse pattern before localizing to puncta towards the apex and cuticular plate at later time points. This is reminiscent of the aminoglycoside loading we observed in zebrafish hair cells, where fluorescently labeled neomycin distributes within hair cells in a diffuse pattern after entry, before becoming sequestered in lysosomes (Hailey et al., 2017). Further characterization of these puncta in mammalian hair cells is needed to better understand their role in aminoglycoside toxicity. Previous in vivo evaluations utilizing GTTR demonstrated the intracochlear trafficking pattern of aminoglycosides, including trans-strial uptake into the endolymph before entering hair cells (Wang and Steyger, 2009). Indeed a recent study demonstrated via real-time imaging using GTTR that aminoglycoside enters the cochlea via the stria vascularis (Kim and Ricci, 2022). Consistent with this analysis, we observe a transient increase in stria vascularis fluorescence before fluorescence returns to near baseline levels by 48 hours post injection. Aminoglycoside uptake in spiral ganglia neurons (SGNs) has previously been observed (Bareggi et al., 1986; Imamura and Adams, 2003; Kitahara et al., 2005), however in our study we did not observe significant fluorescence in the cell bodies of the SGNs. It should, however, be noted that the cumulative dose of aminoglycoside used in these prior in vivo studies was 75-2000 fold higher than the 10 mg/kg administered in this analysis.

An additional extension of this work is the correlation between hair cell aminoglycoside uptake and circulating, systemic levels of aminoglycoside. Clinically, numerous trials have examined the impact of dosing schemes on aminoglycoside toxicity (Barza et al., 1996; Hatala et al., 1996; Munckhof et al., 1996; Rybak et al., 1999; Contopoulos-Ioannidis et al., 2004). Unfortunately, none of these aminoglycoside administration regimens appear to reduce incidence of ototoxicity. In this analysis, we directly correlate plasma levels of G418-TR with G418-TR fluorescence in hair cells and non-sensory tissues of the cochlea. After a single subcutaneous injection of G418-TR, hair cells rapidly accumulate the aminoglycoside at all time points that the G418-TR is detected at any significant level in the serum. While not examined closely in this analysis, the relationship between serum aminoglycoside and hair cell accumulation is likely to be of clinical relevance when considering dosing schemes such as bolus dosing, divided doses, and infusions.

### 4.3 ORC-13661 blocks G418-TR uptake in vivo

Therapeutic inhibition of aminoglycoside uptake is a promising strategy to combat ototoxicity. A recent analysis of gentamicin subtypes demonstrated that chemical modification of the traditional gentamicin mixtures to produce more potent MET channel-blocking analogues can diminishes ototoxicity without compromising antimicrobial activity (O’Sullivan et al., 2020). While aminoglycoside modification remains a promising strategy, multiple previous studies have utilized zebrafish-based screens to identify compounds that inhibit aminoglycoside uptake and protect hair cells (Owens et al., 2008b; Kruger et al., 2016; Kirkwood et al., 2017; Chowdhury et al., 2018; Kitcher et al., 2019; Kenyon et al., 2021). Translating preliminary leads from zebrafish to a mammalian model remains challenging given the resources-intensive techniques required for in vivo mammalian assessment of ototoxicity. Here, we demonstrate that the drug ORC-13661 blocks aminoglycoside uptake *in vivo* in mammals, and we further highlight the utility of the neonatal mouse system to rapidly evaluate ORC-13661 dosing paradigms.

ORC-13661 is a small molecule therapeutic that protects against aminoglycoside ototoxicity in zebrafish and mammals (Chowdhury et al., 2018; Kitcher et al., 2019). Evidence from zebrafish lateral line hair cells and mammalian cochlear cultures suggests that hair cell protection occurs by blocking MET-dependent aminoglycoside uptake (Kitcher et al., 2019). Here, we extend this conclusion to the mammalian inner ear in vivo, demonstrating pharmacologic inhibition of aminoglycoside uptake. We examined two different dosing paradigms and observed a significant dose-dependent reduction in uptake with the short G418-TR exposure paradigm but not the long paradigm. We attribute the observed differences between the two tested schemes to the MET channel block kinetics of ORC-13661. From previous work, we know that ORC-13661 blocks MET-channel currents and MET-dependent aminoglycoside uptake, but this block is readily reversible with washout (Kitcher et al., 2019). Unlike previous in vitro experiments, ORC-13661 is subjected to dynamic in vivo pharmacokinetics in these studies. If the t_1/2_ for ORC-13661 within the plasma or the endolymph of the neonatal mouse is short, the expected therapeutic window following a single injection would also be of limited duration. To test this hypothesis, we developed an ORC-13661 booster treatment scheme. We found that an additional dose of ORC-13661 rescued G418-TR uptake inhibition for animals treated with 30 mg/kg ORC-13661. Our findings suggest that ORC-13661 blocks the uptake of aminoglycosides through reversible inhibition of MET-dependent uptake, and that multiple doses of ORC-13661 may be needed to provide optimal protection from aminoglycosides.

It is worthwhile noting that the doses of ORC-13661 administered in this study (10-100 mg/kg) were larger than those used previously to protect rats (1-5 mg/kg) from aminoglycoside-induced hearing loss (Chowdhury et al., 2018; Kitcher et al., 2019). While this could be due differences between neonatal mice and adult rats, it also raises the question of how much aminoglycoside uptake inhibition is relevant from a hair cell death perspective and a measurable auditory threshold perspective. The association between aminoglycoside uptake and hair cell death is also poorly understood in mammals, in part because the molecular mechanisms of aminoglycoside-induced hair cell death are not completely understood (Forge and Schacht, 2000; Cheng et al., 2005; Rizzi and Hirose, 2007; Jiang et al., 2017). The characteristics of uptake inhibition, and specifically whether level of absolute inhibition, manipulation of rate of uptake, and/or altering the subcellular uptake and localization of aminoglycoside are driving protection are also unknown. Further work to begin correlating aminoglycoside uptake and accumulation with hair cell death and hearing threshold shifts, and conversely aminoglycoside uptake inhibition with hair cell survival and hearing protection is needed to better characterize the therapeutic window for uptake inhibition.

### 4.4 Translational significance

The ability to rapidly evaluate the effects of ototoxins in vivo within the mammalian cochlea will not only enable further characterizations of ototoxins and also novel protective therapeutics. Here, we demonstrate that neonatal mice can treated with the aminoglycoside G418-TR and the kinetics of aminoglycoside uptake can be monitored within the cochlea. We find that cochlear hair cells, especially OHCs, uniquely retain G418-TR for days after a single injection without significant efflux or clearance. Further work is needed to better characterize hair cell retention of aminoglycoside in mammals and correlate aminoglycoside uptake and retention with clinical manifestations of ototoxicity. We further demonstrate that mammalian hair cell uptake of G418-TR can be modulated utilizing ORC-13661, a novel otoprotective therapeutic. These findings support the hypothesis that ORC-13661 protects against aminoglycoside ototoxicity by inhibiting MET-dependent uptake without causing any adverse effect on hearing (Kitcher et al., 2019). Finally, we demonstrate the translational utility of this system by testing a novel ORC-13661 dosing scheme which appears to provide significant and sustained aminoglycoside uptake inhibition. Utilizing the neonatal mouse, we will continue further characterization of additional ototoxic compounds and optimization of protective therapeutic strategies, with the goal of ultimately translating these findings to human clinical trials.

## 5 Author Contributions

JAB, JAS, EWR, and DWR designed the studies. JAB, VR, and LT performed all experiments. AW and JAS provided reagents. JAB and PW acquired data. JAB and DWR analyzed the data and wrote the manuscript.

## 7 Acknowledgements

We are deeply grateful to our laboratory for insightful discussion on the manuscript and T. Linbo for his excellent technical support. This work was supported by AAOHNS ASPO CORE grant (JAB), University of Washington Cystic Fibrosis Foundation Research Development Program fellowship SINGH19R0 (JAB), NIH/NIDCD T32 DC000018 (JAB), and NIH/NIDCD R01 DC005987 (DWR).

