## Supplemental figures for "An in vivo biomarker to characterize ototoxic compounds and novel protective therapeutics"

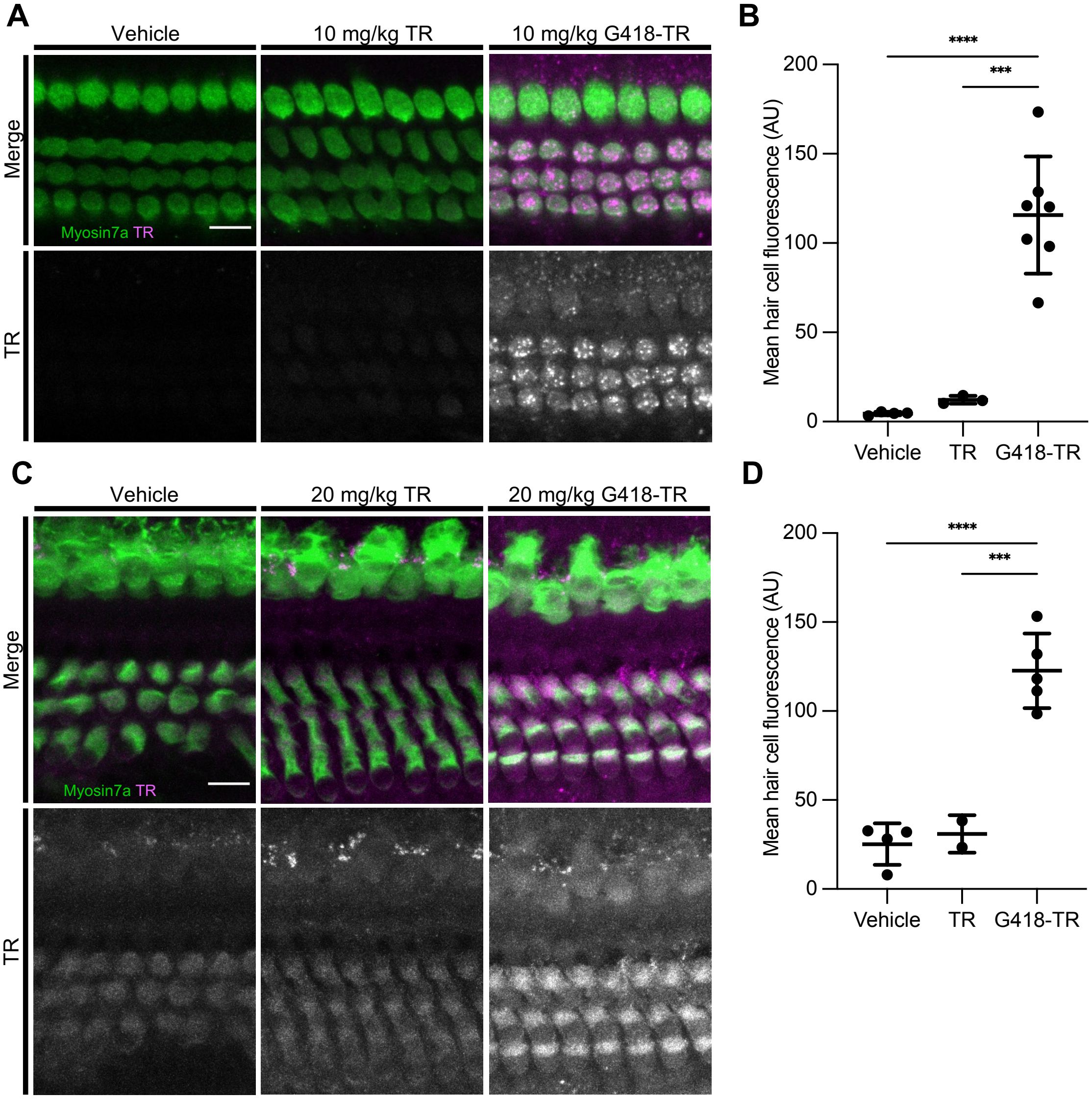


Supplemental Figure 1. **Unconjugated Texas red is not taken up by mammalian hair cells.** (**A**) At 6 hours after systemic injection with vehicle, TR-hydrazide (TR), or G418-TR, neonatal mice (P5) were sacrificed and cochlear sensory epithelia immunolabeled for myosin VIIa. Representative single z-plane images captured from the middle turn of the cochlea in the region between the cuticular plate and nucleus. There is not significant TR uptake observed in hair cells at 6 hours. Scale bar = 10 μm. (**B**) Mean TR fluorescence intensity of neonatal middle turn OHCs 6 hours after systemic treatment. OHCs demonstrate no significant TR uptake when compared to vehicle treated animals (One-way ANOVA with Tukey's multiple comparisons test; P=0.91). (**C**) At 6 hours after systemic injection with vehicle, TR-hydrazide (TR), or G418-TR, juvenile mice (P25-30) were sacrificed and cochlear sensory epithelia immunolabeled for myosin VIIa. Representative single maximum projection images captured from the middle turn of the cochlea. There is not significant TR uptake observed in hair cells at 6 hours. Scale bar = 10 μm. Mean G418-TR fluorescence intensity of IHCs by tonotopic region 6 hours after systemic treatment. IHCs demonstrate similar dose-dependent uptake of G418-TR in base, middle, and apex. (**D**) Mean TR fluorescence intensity of juvenile middle turn OHCs 6 hours after systemic treatment. OHCs demonstrate no significant TR uptake when compared to vehicle treated animals (One-way ANOVA with Tukey's multiple comparisons test; P=0.92). When two cochlea from the same animal were analyzed, the animal mean was calculated and used for all statistics and graphs. Error bars are standard deviations. AU = arbitrary unit. P<0.05, **P<0.01, ***P<0.001, ****P<0.0001; P values generated from multiple comparisons post-hoc test (Tukey's multiple comparison test).

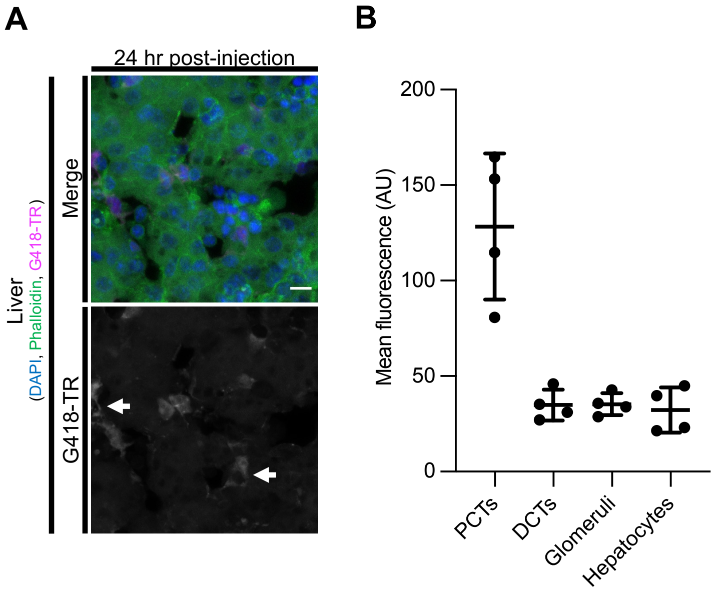


Supplemental Figure 2. **Liver hepatocytes show minimal G418-TR uptake** (**A**) Neonates (P5) treated with 10 mg/kg G418-TR for 24hours show little G418-TR uptake. Note that certain rare cells within the liver (arrow) demonstrate uptake of G418-TR. The morphology of these cells is reminiscent of phagocytic Kupffer cells. Scale bar = 10 µm. (**B**) Quantification of G418-TR uptake 24 hours after injection in the liver and kidney demonstrates low-level uptake in DCTs, Glomeruli, and hepatocytes. Error bars are standard deviations. AU = arbitrary unit and n.s.= not significant.

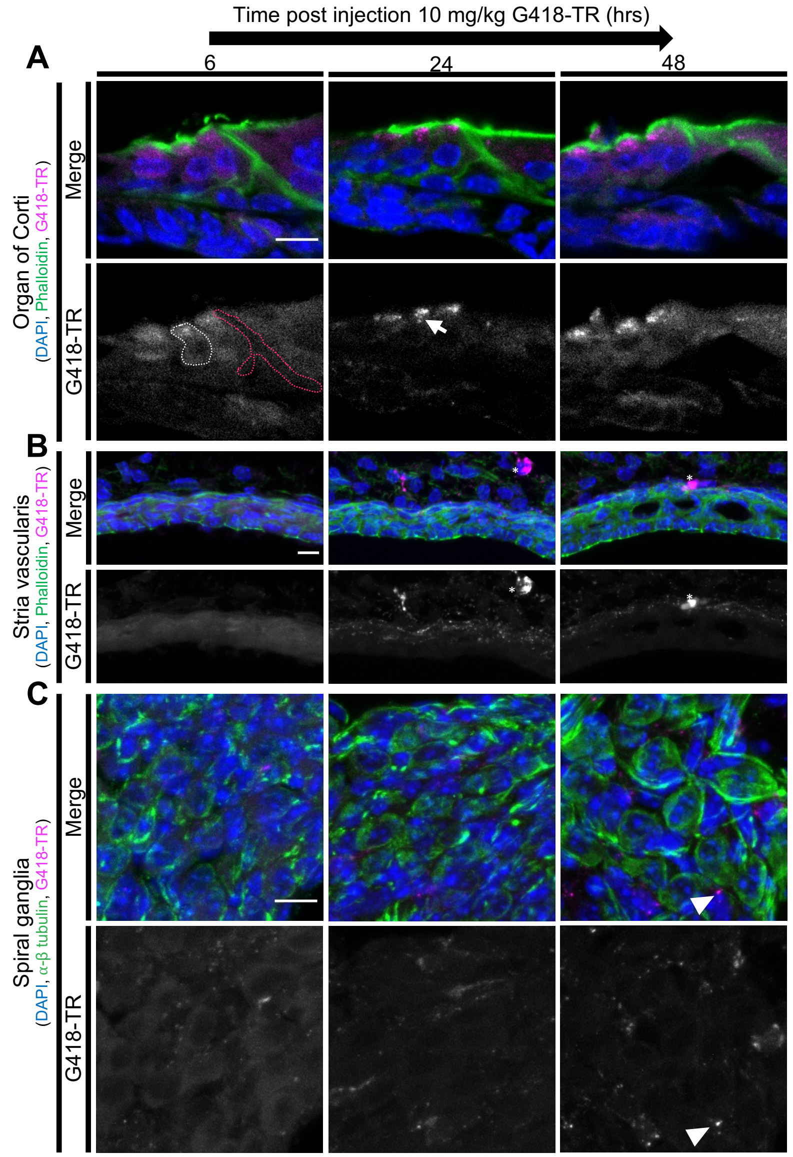


Supplemental Figure 3. **Differential accumulation and retention of G418-TR within the cochlea** (**A**) Cochlea from neonatal mice treated with G418-TR were fixed, cryostat sectioned, and stained with Phalloidin and DAPI. Representative images captured from the middle turn organ of Corti demonstrates G418-TR retention OHCs over time. Apical puncta accumulation of G418-TR (arrow) is again observed. Sensory cells (OHCs, gray dotted lines) and supporting cells (pillar cells, pink dotted line) were specifically analyzed. Scale bar = 10 μm. (**B**) Representative images capture of the Stria vascularis from the middle turn of the cochlea stained with Phalloidin. Stria vascularis tissue takes up G418-TR early, but fluorescence then diminishes over time. Phagocytic cells (asterisk) can be seen accumulating G418-TR in the lateral wall. Scale bar = 10 μm. (**C**) Cryostat sections stained with neuronal class III β-tubulin highlight the spiral ganglia neurons. Minimal uptake is observed in neuronal, however there does appear to be some accumulation in non-neuronal tissue (arrowhead). Scale bar = 10 μm.

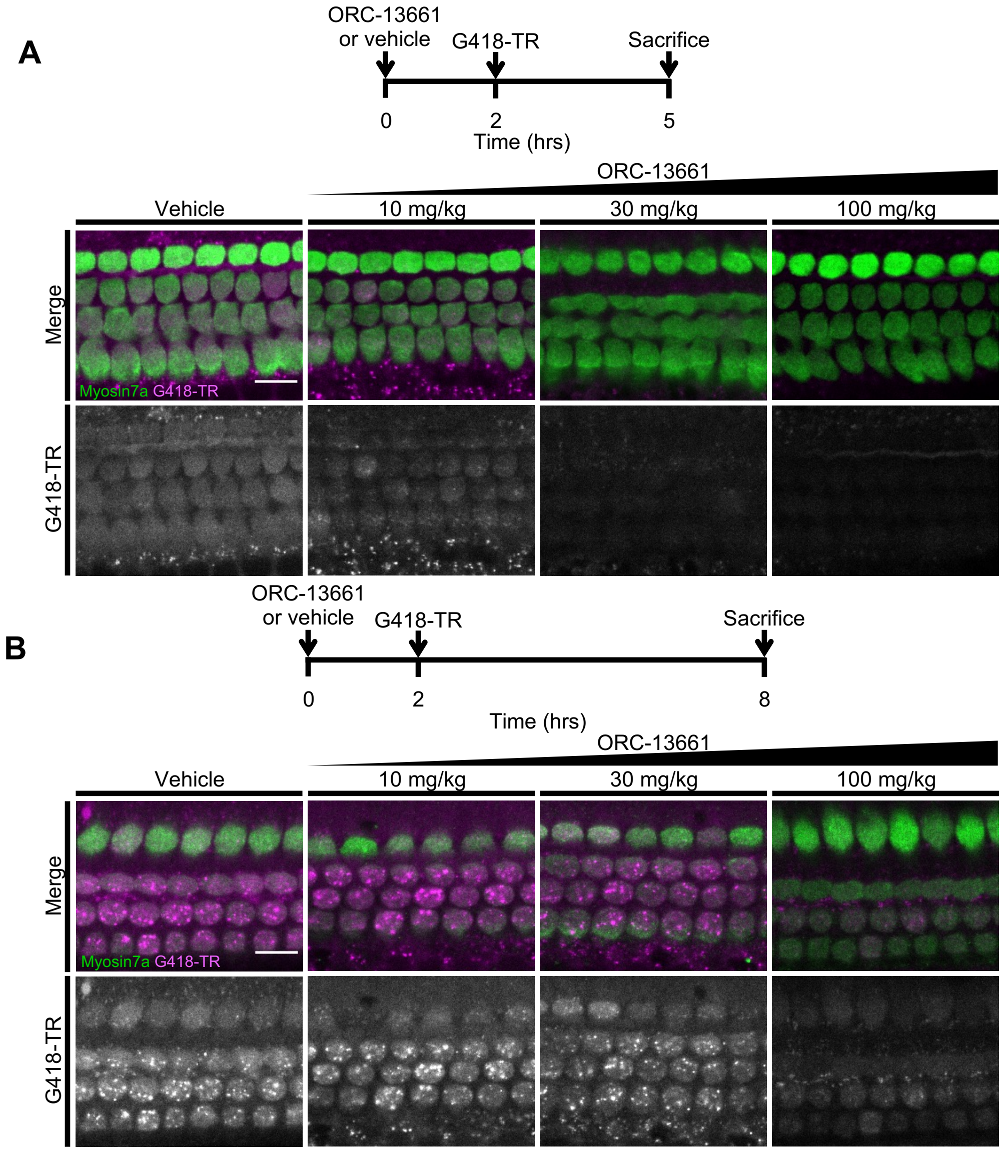


Supplemental Figure 4. **ORC-13661 blocks G418-TR accumulation in mammalian hair cells.** (**A**) Neonatal mice were pretreated with ORC-13661 or vehicle for two hours before injection with 10 mg/kg G418-TR. 3 hours after injection with G418-TR, animals were sacrificed, cochlear sensory epithelium was dissected, fixed, and labeled with myosinVIIa. Representative images from basal turn highlight dose-dependent inhibition of G418-TR uptake with ORC-13661. Scale bar = 10 µm. (**B**) When neonatal mice were pretreated with ORC-13661 or vehicle for two hours, and then exposed to G418-TR for 6 hours, ORC-13661 only blocked uptake at the highest dose. Representative images from basal turn demonstrate significant uptake inhibition with only 100mg/kg ORC-13661. Scale bar = 10 µm.

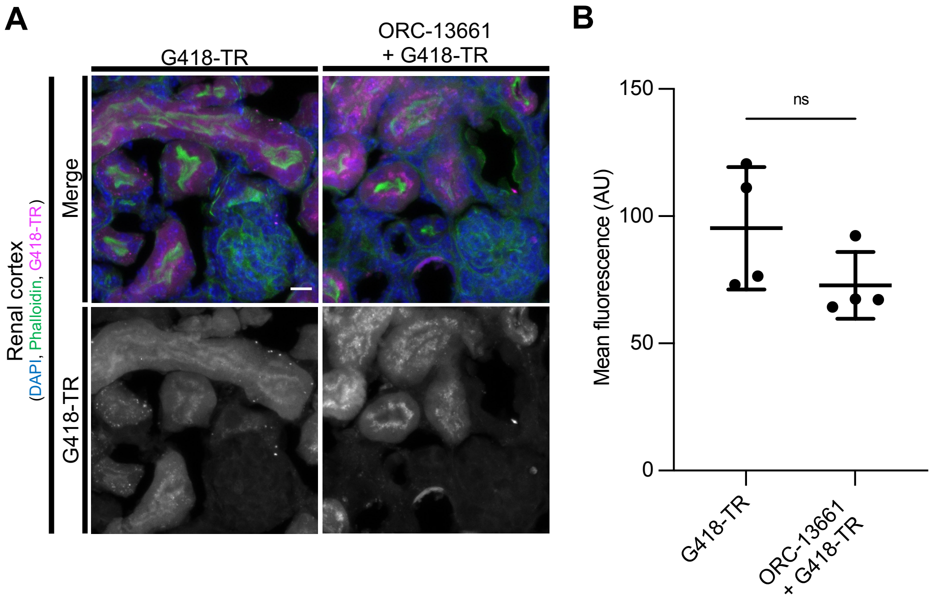


Supplemental Figure 5. **ORC-13661 does not alter G418-TR uptake in the kidney.** (**A**) Neonates (P5) treated with 100 mg/kg ORC-13661 in the 2+6 dosing scheme show similar PCT uptake when compared to animals treated with G418-TR alone. Tissues sections stained with phalloidin and DAPI to highlight the PCT brush border. Note that the pattern of apical accumulation in PCT cells remains present in animals treated with ORC-13661. Scale bar = 10 µm. (**B**) Quantification of PCT G418-TR uptake shows no significant difference between ORC-13661 pretreated animals and those that received G418-TR alone (Unpaired t-test, P=0.34). Error bars are standard deviations. AU = arbitrary unit and n.s.= not significant.
